# A triple fluorescence approach to measure O-GlcNAc dyshomeostasis in stem cells

**DOI:** 10.1101/2025.07.12.664510

**Authors:** Huijie Yuan, Andrew T. Ferenbach, Daan M. F. van Aalten

## Abstract

O-GlcNAcylation is an essential post-translational modification, the complete loss of which results in lethality. Despite modifying thousands of nucleocytoplasmic proteins, O-GlcNAc is controlled by just two enzymes: O-GlcNAc transferase (OGT), which adds the modification, and O-GlcNAcase (OGA), which removes it. Disruptions in O-GlcNAc homeostasis, such as an imbalanced OGT/OGA ratio or aberrant O-GlcNAc levels, are implicated in a wide range of human diseases, including Alzheimer’s disease, cancer, intellectual disability, and diabetes. As such, O-GlcNAc and its regulatory enzymes represent valuable therapeutic targets. However, current tools do not permit informative, large-scale drug or genetic screening, hindering the development of O-GlcNAc-targeted therapies. Here, we present a triple fluorescence stem cell sorting approach in which both endogenous OGT and OGA are tagged with spectrally distinct fluorescent proteins and O-GlcNAc levels can be quantified. We demonstrate that this system faithfully reports disruptions in O-GlcNAc homeostasis. Furthermore, we show that the O-GlcNAc feedback regulation is not solely dependent on O-GlcNAc levels, indicating a role for non-catalytic functions of OGT and OGA. Overall, we provide a high-throughput screening platform that enables reliable and quantitative measurement of O-GlcNAc homeostasis, paving the way for identifying compounds and pathways that target protein O-GlcNAcylation.

## Introduction

O-GlcNAcylation is a non-canonical form of glycosylation that functions as an intracellular post-translational modification (Torres & Hart, 1984). Unlike classical glycosylation, which occurs in the endoplasmic reticulum and Golgi apparatus, O-GlcNAcylation modifies proteins within the cytoplasm, nucleus, and mitochondria. To date, more than 9,000 nucleocytoplasmic proteins have been identified as O-GlcNAc-modified in humans, including protein kinases, transcription factors, RNA-binding proteins, epigenetic regulators, cytoskeletal components, and mitochondrial proteins (Wulff-Fuentes et al., 2021). Given this broad spectrum of substrates, O-GlcNAcylation plays diverse and critical roles in regulating gene expression, RNA splicing, DNA methylation, insulin signaling, immune and stress responses, cell cycle progression, and more (Hart et al., 2007, 2011; Lewis & Hanover, 2014; X. Yang & Qian, 2017; C. Liu & Li, 2018). Not surprisingly, O-GlcNAcylation is essential for life - its complete loss leads to lethality in various metazoan species, including mice and *Drosophila* (Shafi et al., 2000; Sinclair et al., 2009). Remarkably, despite modifying thousands of proteins across diverse pathways and functions, O-GlcNAcylation is regulated by only two enzymes: O-GlcNAc transferase (OGT), the “writer”, which adds the amino sugar UDP-GlcNAc to serine or threonine residues on target proteins (Haltiwanger et al., 1992); and O-GlcNAc hydrolase (OGA), the “eraser”, which removes this modification (Dong & Hart, 1994). OGT and OGA respond to a variety of cellular signals and environmental stimuli, rendering O-GlcNAcylation a dynamic and reversible modification, distinct from canonical glycosylation (Gao et al., 2001; Kreppel & Hart, 1999).

The dynamic nature of O-GlcNAcylation necessitates tight regulatory control to maintain cellular homeostasis. The cycling enzymes OGT and OGA play a central role in this process through a mechanism known as O-GlcNAc feedback regulation. Multiple studies across various cell lines have demonstrated this phenomenon: under high O-GlcNAc conditions, such as those induced by chemical inhibition of OGA, OGT protein and mRNA levels decrease, while OGA protein and mRNA levels increase. Conversely, when O-GlcNAc levels are low, OGT levels rise and OGA levels decline (Slawson et al., 2005; Kazemi et al., 2010; Zhang et al., 2014; Ortiz-Meoz et al., 2015; Burén et al., 2016; Park et al., 2017; Decourcelle et al., 2020; Tan et al., 2021; Yuan et al., 2025; Govindan & Conrad, 2025). This seesaw-like regulation underscores the dynamic nature of the O-GlcNAc system and highlights the importance of maintaining proper O-GlcNAc levels, as disruptions in O-GlcNAcylation are implicated in a wide range of diseases. For instance, reduced O-GlcNAcylation is linked to several neurodegenerative diseases: decreased O-GlcNAcylation of Tau protein has been associated with Alzheimer’s disease (F. Liu et al., 2004), while O-GlcNAc modification of α-synuclein reduces fibril formation, a hallmark of Parkinson’s disease (Marotta et al., 2015). Conversely, elevated O-GlcNAc levels are a common feature of cancer cells, where OGT is frequently overexpressed (Slawson & Hart, 2011). Increased O-GlcNAcylation promotes cell cycle progression and enhances DNA repair mechanisms, contributing to tumorigenesis (Chu et al., 2014; Lewis & Hanover, 2014), while OGT inhibition sensitizes cancer cells to chemotherapy (Caldwell et al., 2010). Similarly, in diabetes, hyperglycemia leads to increased flux through the hexosamine biosynthetic pathway, elevating UDP-GlcNAc levels and, consequently, O-GlcNAcylation, which exacerbates insulin resistance (Marshall et al., 1991). More recently, missense variants in OGT have been identified as the cause of a novel syndromic form of intellectual disability, now classified as OGT-congenital disorder of glycosylation (OGT-CDG) (Vaidyanathan et al., 2017; Willems et al., 2017; Gundogdu et al., 2018; Selvan et al., 2018; Pravata et al., 2019, 2020; Omelková et al., 2023; Mayfield et al., 2024). Collectively, these findings underscore the critical role of O-GlcNAcylation and its cycling enzymes, OGT and OGA, as essential and valuable therapeutic targets across a range of human diseases. Thus, there is a need for tools amenable to large-scale, high-throughput screening, to identify small molecules capable of modulating OGT or OGA activity, uncover genes regulating their protein levels and O-GlcNAc homeostasis, and screen patient-derived OGT and OGA variants to advance personalized therapeutic strategies.

We have previously developed a mouse embryonic stem cell (mESC) line in which the endogenous OGT is fused to the fluorescent marker superfolder GFP (sfGFP), enabling high-throughput screening of potential OGT-CDG variants (Yuan et al., 2025). However, tagging only one of the two O-GlcNAc cycling enzymes may limit the sensitivity and broader applicability of this approach. A more recent study introduced a GFP-based reporter cell line leveraging intron retention as a proxy for O-GlcNAc levels, but as this mechanism operates at the transcriptional level, it is unable to capture post-transcriptional regulation such as translational control or protein degradation (Govindan & Conrad, 2025). Here, we describe a fluorescent mESC line in which both OGT and OGA are endogenously and covalently tagged with spectrally distinct fluorescent proteins—sfGFP and mScarlet3, respectively. We demonstrate that the fluorescence intensities of OGT-sfGFP and mScarlet3-OGA reliably report dynamic changes in cellular O-GlcNAc levels, enabling sensitive and high-throughput detection of disruptions in O-GlcNAc homeostasis. By incorporating a third fluorescent marker for O-GlcNAc immunostaining (RL2-AF647), our system allows simultaneous quantification of endogenous OGT, OGA, and O-GlcNAc levels at the single-cell level. Moreover, we show that the O-GlcNAc feedback regulation is not solely dependent on catalytic activity or O-GlcNAc levels: overexpression of catalytically inactive OGT or OGA variants can still partially trigger the feedback response, suggesting that non-catalytic functions of these enzymes also play a regulatory role, potentially through their extensive interactomes.

## Results and Discussion

### OGT-sfGFP & mScarlet3-OGA double fluorescent mESCs show no apparent changes in O-GlcNAc homeostasis

In continuation of our previously published OGT-sfGFP mESC line, we aimed to engineer the complementary component of the O-GlcNAc cycling system, the O-GlcNAc hydrolase OGA, by tagging it with a spectrally distinct fluorescent reporter. This would generate a dual-fluorescent cell line that enables simultaneous, real-time visualization of both OGT and OGA protein levels. We initially attempted to fuse emiRFP670 either immediately downstream of the last exon or upstream of the first exon of the *Oga* gene in OGT-sfGFP mESCs (Matlashov et al., 2020). However, these approaches resulted in either complete loss of cell viability following antibiotic selection or the emergence of clones with weak fluorescence. Ultimately, we selected a brighter fluorescent protein, mScarlet3, and fused it to the N-terminus of OGA using a flexible eight-amino acid glycine-serine linker (GGGGSGGGGS) via CRISPR gene editing (Fig. 1A) (Gadella et al., 2023). Several single-cell clones were isolated and screened by targeted sequencing. Among these, only one clone exhibited a heterozygous knock-in, with the other allele remaining unmodified and structurally intact (Fig. 1B). In contrast, all other clones displayed varying degrees of genetic disruption in one or both *Oga* alleles. Compared to wild type mESCs and the previously established OGT-sfGFP mESC line, this clone exhibited no significant alterations in either endogenous OGT or OGA protein levels or in global O-GlcNAc levels (Fig. 1B-C). Compared to OGT-sfGFP mESCs, the newly engineered cell line exhibited a marked increase in mScarlet3 fluorescence, while sfGFP fluorescence remained unchanged (Fig. 1D). Notably, immunoblotting with antibodies against OGA and mScarlet3 revealed two closely migrating bands at the expected molecular weight of the mScarlet3-OGA fusion protein (Fig. 1B and 1E). To further investigate this, mScarlet3-OGA was immunoprecipitated using antibodies targeting either OGA or mScarlet3 and probed for both proteins (Fig. 1F). The upper band was consistently enriched, whereas the lower band, though less abundant, may represent a partially degraded form of the mScarlet3-OGA fusion protein. Taken together, these results confirm the successful generation of a dual-fluorescent OGT-sfGFP & mScarlet3-OGA mESC line in which endogenous OGT and OGA are individually tagged with spectrally distinct fluorescent markers, without perturbing O-GlcNAc homeostasis.

**Figure 1:**
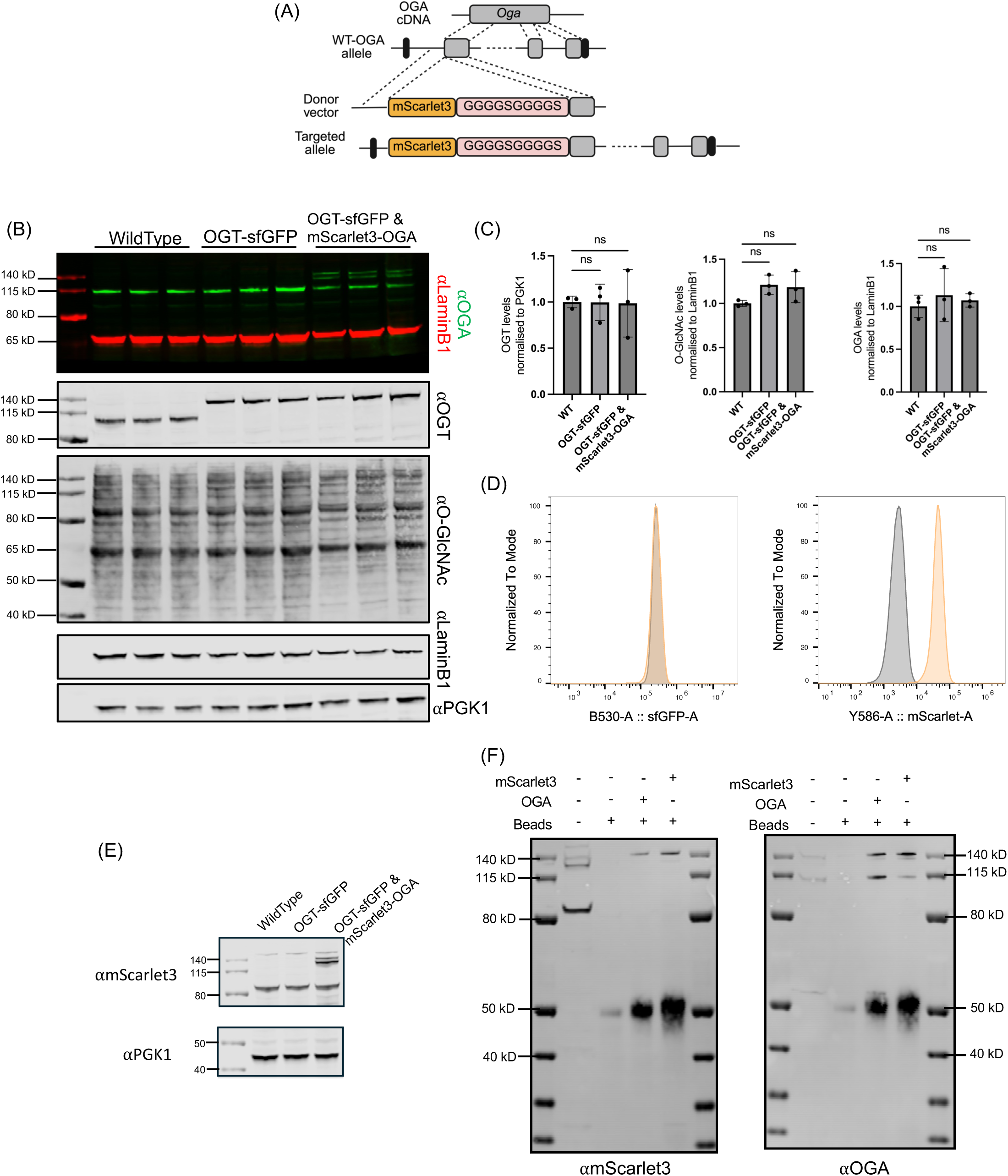
OGT-sfGFP & mScarlet3-OGA double fluorescent mESC line shows no apparent changes in O-GlcNAc homeostasis. **(a)** Schematic representation of CRISPR/Cas9-mediated knock-in of the fluorescent marker mScarlet3 at the N-terminus of OGA using a flexible GS linker (GGGGSGGGGS). **(b)** Immunoblot analysis of cell lysates from unmodified wild-type mESCs, previously reported OGT-sfGFP mESCs, and the newly developed dual-tagged OGT-sfGFP & mScarlet3-OGA mESCs. Experiments were independently performed three times using cells of different passage numbers harvested on separate days. PGK1 and LaminB1 were used as loading controls. Antibody details are provided in the Methods section. **(c)** Quantification of immunoblotting data comparing OGT, OGA, and O-GlcNAc levels across the different genetically modified mESC lines. Statistical analysis was performed using ordinary one-way ANOVA (no matching or pairing). “ns” indicates adjusted *p* values > 0.05. n = 3 independent experiments; error bars represent standard error of the mean (SEM). **(d)** Representative histograms show fluorescence intensity of OGT-sfGFP and mScarlet3-OGA in OGT-sfGFP mESCs and dual-tagged OGT-sfGFP & mScarlet3-OGA mESCs. **(e)** Immunoblotting for mScarlet3 in different fluorescently tagged mESC lines. PGK1 was used as a loading control. **(f)** Immunoprecipitation of OGA or mScarlet3 from dual-tagged OGT-sfGFP & mScarlet3-OGA mESCs, followed by immunoblotting against OGA and mScarlet3.

### Quantification of O-GlcNAc dyshomeostasis through OGT-sfGFP and mScarlet3-OGA fluorescence

O-GlcNAc homeostasis is essential for maintaining proper cellular function, and perturbations in this balance trigger compensatory transcriptional/translational regulation of the two key enzymes in the O-GlcNAc cycling system, OGT and OGA, to minimize disruption (Slawson et al., 2005; Kazemi et al., 2010; Zhang et al., 2014; Ortiz-Meoz et al., 2015; Burén et al., 2016; Park et al., 2017; Decourcelle et al., 2020; Tan et al., 2021; Yuan et al., 2025; Govindan & Conrad, 2025). Leveraging this intrinsic feedback mechanism, the protein levels of OGT and OGA can serve as sensitive readouts for disturbances in O-GlcNAc homeostasis. To assess whether mScarlet3-OGA fluorescence faithfully reflects endogenous OGA protein levels, the dual-tagged OGT-sfGFP & mScarlet3-OGA mESCs were treated with the OGT inhibitor OSMI-4 and the OGA inhibitor Thiamet-G (TMG) for 24 h. Immunoblotting revealed evidence for this feedback regulation: OSMI-4 treatment reduced global O-GlcNAc levels, resulting in elevated OGT and markedly diminished OGA protein levels, whereas TMG treatment increased O-GlcNAc levels, leading to elevated OGA and reduced OGT protein levels (Fig. 2A).

**Figure 2:**
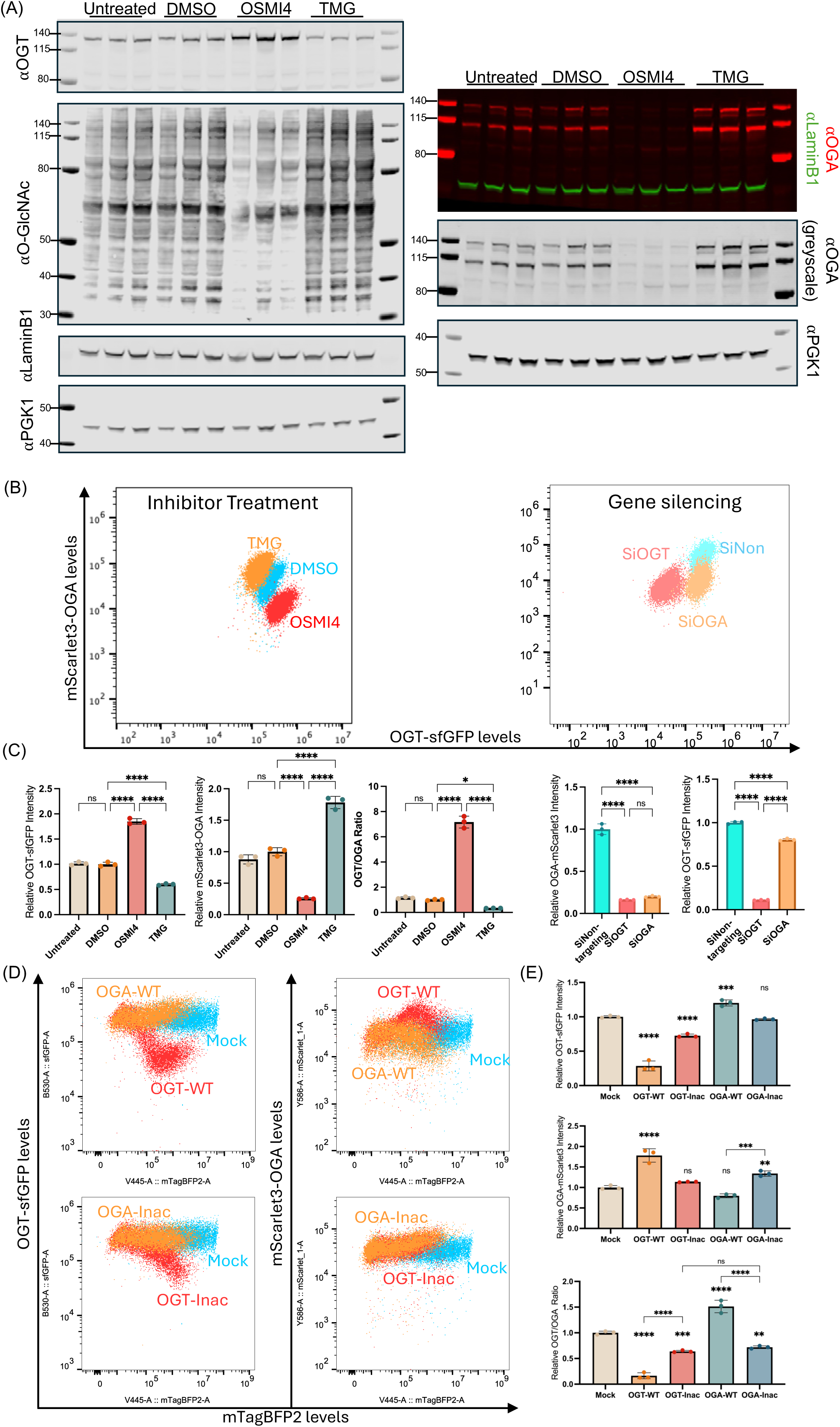
OGT-sfGFP & mScarlet3-OGA fluorescence probes disruptions in O-GlcNAc homeostasis. **(a)** Immunoblotting of OGT, OGA, and O-GlcNAc levels from lysates of OGT-sfGFP & mScarlet3-OGA mESCs treated with 10 µM OSMI-4 or 10 µM TMG for 24 h. Cells treated with 0.1% DMSO served as controls. PGK1 and LaminB1 were used as loading controls. **(b)** Density plots of OGT-sfGFP (x-axis) and mScarlet3-OGA (y-axis) fluorescence following different treatments. The left panel shows responses to 24 h treatment with 10 µM OSMI-4 or TMG. The right panel shows effects of siRNA-mediated knockdown of either OGT or OGA. **(c)** Quantification of OGT-sfGFP fluorescence, mScarlet3-OGA fluorescence, and OGT/OGA fluorescence ratios following treatments shown in panel (b). Statistical analysis was performed using ordinary one-way ANOVA (n = 3 independent replicates). Error bars represent mean ± SEM. Significance: ∗ for adjusted *p* < 0.05, ∗∗∗∗ for *p* < 0.0001, and ns for adjusted *p* > 0.05. **(d)** Density plots show OGT-sfGFP and mScarlet3-OGA fluorescence in relation to mTagBFP2 fluorescence in cells transfected with mTagBFP2-labeled constructs expressing either wild-type (OGT-WT/OGA-WT) or catalytically inactive (OGT-Inac/OGA-Inac) variants of OGT or OGA. A plasmid encoding only the mTagBFP2 marker (Mock) was used as a control. Each dot represents a single cell; colors and labels indicate transfected construct. **(e)** Quantification of endogenous OGT-sfGFP and mScarlet3-OGA fluorescence intensity and OGT/OGA fluorescence ratios in mTagBFP2-positive cells transfected with the constructs shown in panel (d). Statistical analysis was performed using ordinary one-way ANOVA (n = 3 independent replicates). Error bars represent mean ± SEM. Significance: ∗∗ for adjusted *p* < 0.01, ∗∗∗ for *p* < 0.001, ∗∗∗∗ for *p* < 0.0001, and ns for adjusted *p* > 0.05. Significance levels were indicated above each histogram relative to the mock transfected controls; additional pairwise comparisons were labeled accordingly.

Flow cytometry analysis of cells subjected to the same treatments revealed distinct separation between OSMI-4 and TMG treated populations based on sfGFP and mScarlet3 fluorescence (Fig. 2B, left panel). Quantification of fluorescence intensities mirrored the immunoblotting results, demonstrating the fidelity of the dual-fluorescent reporters in capturing the feedback regulation of OGT and OGA (Fig. 2C, left three graphs). These findings demonstrate a robust and high-throughput approach to simultaneously monitor endogenous OGT and OGA protein levels and to detect perturbations in O-GlcNAc homeostasis by measuring sfGFP and mScarlet3 fluorescence. Notably, the ratio of OGT to OGA fluorescence can serve as an informative parameter for quantifying the extent of homeostatic disruption (Fig. 2C).

The dual-tagged OGT-sfGFP & mScarlet3-OGA mESCs were also subjected to siRNA-mediated knockdown of OGT (siOGT) and OGA (siOGA) to further investigate feedback regulation within the O-GlcNAc system. Interestingly, these perturbations revealed a non-reciprocal feedback response, resulting in separation of cell populations across different treatments (Fig. 2B, right panel). Specifically, siOGT treatment led to a reduction in OGT-sfGFP fluorescence, accompanied by a concomitant decrease in mScarlet3-OGA fluorescence, consistent with a coordinated feedback mechanism (Fig. 2C, right two graphs). However, siOGA treatment significantly reduced mScarlet3-OGA fluorescence, but had less effect on OGT-sfGFP levels (Fig. 2C, right two graphs). These findings suggest that OGT and OGA are subject to feedback regulation through mechanistically distinct, and potentially asymmetric, pathways.

We previously demonstrated that OGT-sfGFP fluorescence responds to the overall activity of transfected exogenous OGT variants, providing a high-throughput platform to screen potential OGT-CDG variants (Yuan et al., 2025). To assess whether the mScarlet3-OGA fusion protein behaves similarly, we transfected the dual-tagged OGT-sfGFP & mScarlet3-OGA mESCs with plasmids encoding wild-type OGT (OGT-WT) and OGA (OGA-WT), as well as their catalytically inactive counterparts: OGT-Inac (bearing K842M/H901Y mutations) and OGA-Inac (bearing D174A/D175N mutations). All constructs were linked to a third fluorescent marker, mTagBFP2, through a P2A linker, enabling the selective analysis of transfected mTagBFP2 positive cells, as previously described (Yuan et al., 2025) and illustrated in the gating strategy (Supplementary Figure 1). A plasmid encoding only the mTagBFP2 marker (Mock) was used as a control. As expected, the resulting density plots following plasmid transfection illustrate how OGT-sfGFP and mScarlet3-OGA levels respond to increasing expression of the exogenous variants (Fig. 2D, left panel). Upon OGT-WT transfection, a rise in mTagBFP2 fluorescence, which indicates increased transgene expression, correlated with a decrease in OGT-sfGFP fluorescence, corresponding to an approximate 2/3 reduction compared to the Mock control. This was accompanied by a corresponding increase in mScarlet3-OGA fluorescence, reaching about 1.5 times the baseline mScarlet3 level observed in Mock control (Fig. 2D). In contrast, OGA-WT transfection elicited a less significant response, with OGT-sfGFP fluorescence increasing by roughly 1.2-fold and mScarlet3-OGA fluorescence showing a not statistically significant decrease (Fig. 2D).

Interestingly, transfection with the catalytically inactive OGT-Inac mutant also resulted in a reduction of OGT-sfGFP fluorescence, albeit to a lesser extent than OGT-WT. The sfGFP fluorescence intensity decreased to approximately 80% of the baseline observed in the Mock control, without significantly affecting mScarlet3-OGA levels (Fig. 2D). In contrast, OGA-Inac transfection did not alter OGT-sfGFP fluorescence but led to an increase, approximately 1.2-fold, in mScarlet3-OGA intensity. Collectively, these findings confirm that in the dual-tagged OGT-sfGFP & mScarlet3-OGA mESC line, sfGFP and mScarlet3 fluorescence reliably reflect endogenous OGT and OGA protein levels, thereby serving as dynamic reporters of O-GlcNAc homeostasis. Furthermore, the asymmetric feedback responses observed between OGT and OGA suggest that the regulatory mechanisms governing their expression are, at least in part, distinct.

### O-GlcNAc feedback regulation is not solely O-GlcNAc dependent

The observation that catalytically inactive OGT and OGA mutants can partially elicit feedback responses raises the possibility that these mutants may modulate global O-GlcNAc levels through non-catalytic mechanisms in a cellular context. Both OGT and OGA are known to form homodimers, and dimerization is critical for OGA’s catalytic function (Jinek et al., 2004; B. Li et al., 2017). It is therefore plausible that overexpression of catalytically inactive OGA interferes with endogenous OGA activity by forming heterodimers with wild type OGA, thereby reducing the overall enzymatic activity of the complex. To investigate this, we employed a fluorescently conjugated anti-O-GlcNAc antibody (RL2-AF647) to quantify cellular O-GlcNAc levels following fixation and permeabilization. First, we assessed whether the fixation and staining procedure interferes with endogenous sfGFP and mScarlet3 fluorescence. Dual-tagged OGT-sfGFP & mScarlet3-OGA mESCs were treated with either OSMI-4 or TMG, fixed, and stained with RL2-AF647. Fixation and staining had no measurable impact on sfGFP or mScarlet3 fluorescence, and the expected changes in global O-GlcNAc levels were observed: OSMI-4 treatment decreased RL2-AF647 fluorescence, while TMG treatment increased it (Fig. 3A). Next, cells were transfected with plasmids encoding OGT-WT, OGA-WT, or the mock construct expressing only the mTagBFP2 marker. After fixation and RL2-AF647 staining, density plots revealed that cell size decreased upon fixation, while mTagBFP2 fluorescence remained unaffected (Fig. 3B). Within the mTagBFP2 positive population, the endogenous sfGFP and mScarlet3 signals remained consistent with previous observations, confirming that the fixation and staining procedures had minimal impact upon endogenous OGT-sfGFP or mScarlet3-OGA fluorescence (Fig. 3C). As expected, expression of OGT-WT led to elevated global O-GlcNAc levels, whereas OGA-WT expression resulted in reduced O-GlcNAc levels (Fig. 3C). Together, these results show that OGT-sfGFP and mScarlet3-OGA fluorescence serve as indirect reporters of global O-GlcNAc levels. When combined with a third fluorescent marker, RL2-AF647, this system enables a high-throughput platform for interrogating O-GlcNAc homeostasis and its perturbations at both the protein expression and modification levels.

**Figure 3:**
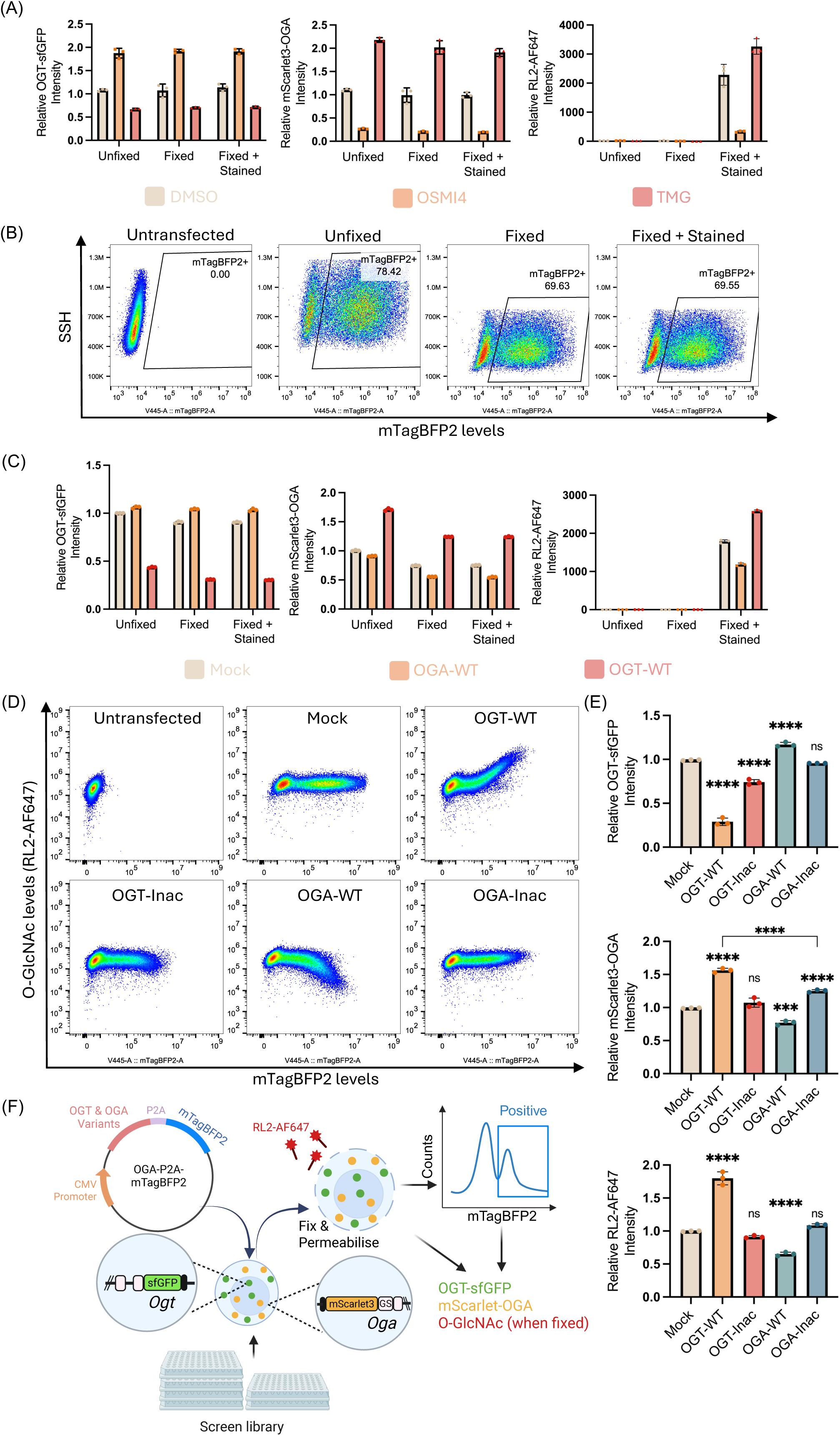
O-GlcNAc feedback regulation is not exclusively O-GlcNAc dependent. **(a)** Quantification of OGT-sfGFP, mScarlet3-OGA, and RL2-AF647 fluorescence intensities in dual-tagged mESCs treated with 10 µM OSMI-4 or TMG. Cells were either directly subjected to flow cytometry (Unfixed), fixed and permeabilized (Fixed), or fixed, permeabilized, and stained with RL2-AF647 (Fixed + Stained). Cells from the same passage were used for all treatments, with each condition comprising three technique replicates. Cells were harvested and analyzed on the same day. Error bars represent mean ± SEM. **(b)** Representative density plots show mTagBFP2 (x-axis) and side scatter height (y-axis) in dual-tagged mESCs transfected with mock (mTagBFP2-only) plasmid. An untransfected control was used to define gating for mTagBFP2-positive cells. Transfected cells were analyzed under the same three conditions as in panel (a). **(c)** Quantification of OGT-sfGFP, mScarlet3-OGA, and RL2-AF647 fluorescence intensities in mTagBFP2 positive cells transfected with mock plasmid, OGT-WT, or OGA-WT constructs. Cells were analyzed under the same three conditions as above. Each data point represents an individual replicate; cells were harvested and analyzed on the same day with error bars represent mean ± SEM. **(d)** Density plot shows mTagBFP2 (x-axis) versus RL2-AF647 (y-axis) fluorescence in dual-tagged mESCs transfected with either mock plasmid, OGT-WT, OGA-WT, catalytically inactive OGT (OGT-Inac), or catalytically inactive OGA (OGA-Inac). All samples were fixed and stained with RL2-AF647. Each dot represents a single cell. **(e)** Quantitative analysis of OGT-sfGFP, mScarlet3-OGA, and RL2-AF647 fluorescence in mTagBFP2 positive cells shown in panel (d). Statistical analysis was performed using ordinary one-way ANOVA (n = 3 independent replicates). Error bars indicate mean ± SEM. Significance: ∗∗∗ for adjusted *p* < 0.001, ∗∗∗∗ for adjusted *p* < 0.0001, and ns for adjusted *p* > 0.05. Significance values are shown above each histogram relative to the Mock control; additional pairwise comparisons are labelled directly. **(f)** Schematic illustration of the high-throughput O-GlcNAc dyshomeostasis screening platform reported in this work. Dual-tagged OGT-sfGFP & mScarlet3-OGA mESCs can be transfected with mTagBFP2-labeled OGT/OGA variants or exposed to chemical libraries. Following treatment, cells are fixed, permeabilized, stained with RL2-AF647, and analyzed by flow cytometry. For transfection screens, mTagBFP2 positive cells are gated prior to downstream analysis. For chemical screens, the entire cell population is assessed. Fluorescence intensities of OGT-sfGFP, mScarlet3-OGA, and RL2-AF647 are used to quantify changes in O-GlcNAc homeostasis.

Using this approach, catalytically inactive OGT and OGA mutants were transfected into the dual-fluorescent reporter cells, followed by fixation and RL2-AF647 staining. As shown in the density plots, overexpression of OGT-Inac and OGA-Inac did not alter global O-GlcNAc levels, similar to the mock control (Fig. 3D). However, consistent with previous observations in unfixed cells, OGT-Inac expression led to a reduction in OGT-sfGFP fluorescence, while OGA-Inac expression resulted in increased mScarlet3-OGA fluorescence (Fig. 3E). These findings suggest that O-GlcNAc feedback regulation is not only dependent on the catalytic activity of OGT or OGA. Non-catalytic functions, such as protein–protein interactions, may also contribute to this regulation, albeit more subtly. Overall, the data demonstrate that O-GlcNAc feedback regulation is not solely O-GlcNAc dependent.

## Concluding remarks

Here, we engineered a dual-fluorescent mESC line in which the two key enzymes regulating O-GlcNAcylation, the writer OGT and the eraser OGA, are covalently fused to spectrally distinct fluorescent proteins. Attempts to tag OGA by directly fusing a ∼25 kDa fluorescent protein to either its N-or C-terminus failed to yield fluorescent clones. However, successful tagging was achieved by fusing the brighter fluorescent protein mScarlet3 to the N-terminus of OGA via a flexible GS linker. Human full-length OGA is a multidomain protein composed of an N-terminal O-GlcNAc hydrolase domain, a central stalk linker domain, and a C-terminal pseudo-histone acetyltransferase (pHAT) domain (Roth et al., 2017). The failure of direct N-or C-terminal tagging suggests that such fusions may interfere with OGA function or stability, possibly mimicking the effects observed in OGA loss-of-function lethality (Keembiyehetty et al., 2015; Muha et al., 2021; Y. R. Yang et al., 2012). Notably, the pHAT domain has been shown to be essential for recruiting OGA to DNA lesions (Cui et al., 2021), suggesting that direct fusion at this site may disrupt key protein interactions. We successfully isolated only one heterozygously tagged mScarlet3-OGA clone, with the second allele remaining intact and unmodified. Notably, this heterozygous tagging did not result in significant alterations in endogenous OGT, OGA, or global O-GlcNAc levels. Interestingly, immunoblotting of cell lysates revealed two closely migrating bands (∼140 kDa) of the mScarlet3-OGA fusion protein. The upper band was consistently enriched during immunoprecipitation with either anti-OGA or anti-mScarlet3 antibodies. These bands are unlikely to represent the short isoform of OGA (Comtesse et al., 2001; E. J. Kim et al., 2006), which typically migrates around 75 kDa. Instead, the observed doublet likely reflects either partial proteolytic truncation of the fusion protein or distinct conformational states that affect its electrophoretic mobility.

By labelling OGT and OGA with spectrally different fluorescent markers, we demonstrate that OGT-sfGFP and mScarlet3-OGA fluorescence reflects their respective protein levels. These signals respond to pharmacological inhibition or gene silencing of OGT or OGA, establishing this system as a dual fluorescent reporter for monitoring disruptions in O-GlcNAc homeostasis. This enables high-throughput screening of small molecules or genetic perturbations that modulate O-GlcNAc levels in specific directions. For instance, compounds that reduce OGT-sfGFP fluorescence and increase mScarlet3-OGA fluorescence likely elevate global O-GlcNAc levels, offering potential for therapeutic development in OGT-CDG and other hypo-O-GlcNAc related neurodegenerative diseases. Conversely, compounds that increase OGT-sfGFP and decrease mScarlet3-OGA fluorescence likely reduce O-GlcNAc levels, which may be beneficial for targeting diseases such as diabetes and cancer, where O-GlcNAcylation is often abnormally elevated. However, it is important to note that OGT and OGA protein levels are also regulated by factors beyond the feedback mechanism (Z. Li et al., 2017, p. 1; C. Liu & Li, 2018; Muthusamy et al., 2015; Zachara et al., 2004). Since OGT-sfGFP and mScarlet3-OGA fluorescence are indirect measures of O-GlcNAc levels, validating any observed effects requires direct assessment of cellular O-GlcNAc levels.

Using a third fluorescent probe, RL2-AF647, we established a method to fix and stain dual fluorescently tagged OGT-sfGFP & mScarlet3-OGA mESCs for O-GlcNAc detection. Importantly, our fixation, permeabilization, and antibody staining protocol preserves the endogenous sfGFP and mScarlet3 fluorescence, enabling simultaneous measurement of OGT, OGA, and global O-GlcNAc levels within the same cells. To assess the impact of exogenous proteins, constructs can be co-expressed with a fourth fluorescent marker, mTagBFP2, linked via the self-cleaving P2A peptide (J. H. Kim et al., 2011). This enables identification of transfected cells, even after fixation, by gating on mTagBFP2 fluorescence. Together, this system constitutes a multiplexed, tetra-fluorescent platform for probing the entire O-GlcNAc axis (OGT, OGA, and O-GlcNAc) in a high-throughput and quantitative manner. Its sensitivity and scalability make it well-suited for downstream applications such as small-molecule screening, genome-wide CRISPR/Cas9 perturbation studies, and broader investigations into the regulation of O-GlcNAc homeostasis. With the growing number of OGT variants linked to human diseases such as intellectual disability (Mayfield et al., 2024), the tetra-fluorescent system offers a rapid and comprehensive approach to assess how different OGT, and potentially OGA variants impact O-GlcNAc homeostasis, thereby aiding in the prediction of disease-associated variants.

While our method relies on the O-GlcNAc feedback regulation, the underlying mechanisms remain poorly understood. Several studies have described how OGT and OGA protein levels are modulated under conditions such as cell cycle progression (C. Liu & Li, 2018), cancer (Caldwell et al., 2010) and stress response (M. R. Martinez et al., 2017); however, the extent to which these mechanisms contribute to the O-GlcNAc feedback regulation remains unclear. Both OGT and OGA genes contain a highly conserved intronic region that mediates intron retention, leading to nuclear detention of transcripts containing them (Park et al., 2017; Tan et al., 2021). This provides a potential explanation for feedback regulation at the transcriptional level. For example, within just 2 h of OGT inhibitor treatment, there is a reduction in the levels of intron-retaining OGT and OGA transcripts (Tan et al., 2021). This correlates with an increase in functional OGT transcripts and non-functional OGA transcripts, along with a decrease in functional OGA transcripts and non-functional OGT transcripts (Tan et al., 2021). A recent study further implicated the splicing factor SFSWAP in regulating OGT intron retention (Govindan & Conrad, 2025). However, SFSWAP appears to function as a more general regulator of intron retention, rather than being specific to OGT or the O-GlcNAc system. While these findings shed light on transcriptional-level feedback, the mechanisms underlying translational and post-translational regulation of OGT and OGA in this context remain largely unknown. We show that the feedback regulation of OGT and OGA is not reciprocal. Knockdown of OGT results in a significant reduction of OGA levels, whereas knockdown of OGA leads to a smaller decrease in OGT. Moreover, overexpression of catalytically inactive OGT or OGA constructs, while not altering global O-GlcNAc levels, is still sufficient to partially trigger feedback regulation: overexpression of inactive OGA increases endogenous OGA levels, and inactive OGT decreases endogenous OGT levels. These results indicate that the feedback mechanism is not solely dependent on O-GlcNAc levels. Rather, the non-catalytic functions of OGT and OGA can also contribute to feedback regulation, albeit to a lesser extent than their wild-type counterparts. This may be linked to the extensive interactomes of both enzymes (M. Martinez et al., 2021). Further investigation is needed to elucidate the underlying mechanisms. Overall, this study presents a multiplexed tetra-fluorescent reporter system that enables quantitative assessment of disruptions in O-GlcNAc homeostasis. This system offers a powerful platform for large-scale, high-throughput screening and a valuable tool for dissecting the molecular basis of O-GlcNAc feedback regulation.

## Materials & Methods

### CRISPR mScarlet3-OGA knock-in

OGT-sfGFP mESCs were maintained in GMEM-BHK 21 medium (Gibco, 11710035) supplemented with 10% [v/v] fetal bovine serum (Gibco, A5256701), 1 mM sodium pyruvate (Gibco, 11360088), 1000 U/mL LIF (Merck, ESG1107), 0.1 mM MEM non-essential amino acids (Gibco, 11140050), and 0.1 mM 2-mercaptoethanol (Gibco, 31350010). Cells were transfected with Cas9 D10A nickase, paired guide RNAs, and a repair template using Lipofectamine 3000, according to the manufacturer’s protocol. Following transfection, puromycin selection (1 µg/mL) was applied for 48 h. Successful genomic integration was verified by restriction enzyme digestion and Sanger sequencing of genomic DNA. Details of CRISPR/Cas reagents and cloning procedures are available in our previously published study (Yuan et al., 2025).

### Immunoblotting

Cells were harvested, washed with PBS, and lysed in RIPA buffer (Thermo Fisher). Protein concentrations were determined using the Pierce™ 660 nm Protein Assay Reagent (Thermo Fisher). Equal amounts of crude lysate (20 µg) were resolved on 4–12% Bis-Tris NuPAGE gels (Invitrogen) and transferred to nitrocellulose membranes using a semi-dry transfer system. Membranes were blocked in 5% [w/v] BSA in TBS-T for at least 1 h at room temperature, followed by incubation with primary antibodies under the specified conditions (see below). After three 5 min washes in TBS-T, membranes were incubated with IRDye 680RD or IRDye 800CW secondary antibodies (LI-COR) for 1 h at room temperature. Blots were imaged using the Odyssey CLx system (LI-COR) with 700 nm and 800 nm detection channels.

The primary antibodies used were: OGT (Sigma, DM-17; 1:1000; 2 h, room temperature), RL2 for O-GlcNAc (Thermo Fisher; 1:500; overnight at 4°C), LaminB1 (Proteintech, 12987-1-AP; 1:5000; 1.5 h, room temperature), PGK1 (Proteintech, 17811-1-AP; 1:5000; 1.5 h, room temperature), OGA (Thermo Fisher, SAB4200267; 1:1000; 4 h, room temperature), and pan-RFP for mScarlet3 (Chromotek, pabr1; 1:5000; 1.5 h, room temperature).

### Immunoprecipitation

A total of 200 µg of protein lysate from dual-fluorescent OGT-sfGFP & mScarlet3-OGA mESCs was incubated overnight at 4 °C with either 1.5 µg anti-OGA antibody or 3 µg anti-mScarlet3 (pan-RFP). The antibody-lysate mixtures were then added to pre-washed Dynabeads™ Protein A (Invitrogen™) and incubated for 10 min at room temperature with gentle mixing. Beads were subsequently washed three times with 200 µl wash buffer (PBS supplemented with 0.02% Tween-20). For elution, 20 µl of 50 mM glycine (pH 2.8) was added to the beads and incubated for 5 min at room temperature with rotation. The eluates were neutralized by adding 2.5 µl of 1 M Tris-HCl (pH 8.0), followed by addition of 7.5 µl 4× LDS sample buffer (Invitrogen) containing TCEP. Samples were then boiled for 5 min at 95 °C and the supernatant collected for western blot analysis.

### OSMI-4b and Thiamet G (TMG) Treatment

For immunoblotting, 0.5 million dual-fluorescent OGT-sfGFP & mScarlet3-OGA mESCs were seeded per well in 6-well plates. After 24 h, cells were treated with 10 µM OSMI-4b or 10 µM Thiamet G (TMG), each prepared in 0.1% DMSO. A 0.1% DMSO vehicle control and a no-treatment control were included. After 24 h treatment, cells were harvested for immunoblotting. For flow cytometry, 0.1 million dual-fluorescent mESCs were seeded per well in 12-well plates and treated under the same conditions. Following 24 h of drug exposure, cells were harvested and analyzed by flow cytometry (see gating strategy in Supplementary Figure 1). The median sfGFP and mScarlet3 fluorescence intensities were extracted for statistical analysis. Both experiments were independently repeated three times on different days using cells from different passage numbers. Stock solutions of OSMI-4b and TMG were prepared in 100% DMSO at a concentration of 10 mM.

### Transfection

For all transfection experiments, OGT-sfGFP mESCs were cultured in 2i medium to enhance transfection efficiency (Ying et al., 2008). The 2i medium consisted of 50% DMEM/F-12 with GlutaMAX™ Supplement (Gibco, 31331-093) and 50% Neurobasal (Gibco, 21103049), supplemented with 1×N-2 Supplement (Gibco, 17502048), 1×B27 Supplement (Gibco, 17504044), 1 µM MEK inhibitor PD0325901 (Merck, PZ0162), 3 µM GSK3 inhibitor CHIR99021 (Merck, SML1046), 1000 U/ml LIF (Merck, ESG1107), and 2% [v/v] FBS (Gibco, A5256701). Transfections were performed following a previously published protocol (Tamm et al., 2016). Briefly, 1.5 µg plasmid DNA was mixed with 3 µl Lipofectamine 2000 (Thermo Fisher) in 100 µl Opti-MEM (Thermo Fisher) according to the manufacturer’s instructions. A suspension of 3.0 × 10⁵ cells was incubated with the transfection complex for 5 min prior to seeding into 12-well plates. After 24 h, the medium was replaced, and cells were harvested for flow cytometry after an additional 24 h. This procedure was used for all transfection experiments not involving fixation. For experiments followed by fixation, 5.0 × 10⁵ cells were used instead.

### Fixation and RL2-Alexfluor 647 staining

The harvested cells were fixed in warm 4% paraformaldehyde (PFA) with 4% sucrose for 10 min and washed with PBS. Nonspecific binding was blocked and the cells were permeabilized using blocking solution (5 % BSA and 0.3 % Triton X-100 in PBS) for 1 h at room temperature. Cells were then incubated with 2 µl RL2-AlexaFluor647 in 100 µl diluted blocking solution (1 % BSA in PBS) overnight at 4 °C. After washing with PBS, cells were resuspended in PBS for flow cytometry analysis. For unfixed cell samples, cells were resuspended in PBS supplemented with 1X RedDot™2 live/dead dye (Biotium) and immediately analysed by flow cytometry.

### Flow cytometry analysis

For sample analysis, a NovoCyte Quanteon 4025 flow cytometer (Agilent, Santa Clara, CA) equipped with four lasers (405 nm, 488 nm, 561 nm, and 637 nm) and 25 fluorescence detectors was used. mTagBFP2 fluorescence was detected using the 405 nm laser and a 445/45 bandpass filter. The 488 nm laser combined with a 561/14 filter was used for forward scatter (FSC), while the 561 nm laser and 561/14 filter were used for side scatter (SSC). sfGFP fluorescence was detected with the 488 nm laser and a 530/30 filter. mScarlet3 fluorescence was measured using the 561 nm laser and a 586/20 filter. RL2-AlexaFluor647 (AF647) was detected using the 637 nm laser and a 667/30 filter. RedDot2 was detected using the 637 nm laser and a 685-705 nm filter. Data acquisition was performed using the NovoExpress software (version 1.6.2), and analysis was conducted using FlowJo™ (version 10). The gating strategy is detailed in Supplementary Figure 1.

### Statistical analysis

All statistical analyses were performed using GraphPad Prism (version 10). Ordinary one-way ANOVA without matching or pairing was applied, and adjusted *p* values are indicated on the corresponding graphs.

## Acknowledgements

This work was funded by a Wellcome Trust Investigator Award (110061), a Novo Nordisk Fonden Laureate award (NNF21OC0065969) and a Villum Fonden Investigator (00054496) to D.M.F.v.A. Supported in part by the Danish Research Institute of Translational Neuroscience – DANDRITE of the Nordic-EMBL Partnership for Molecular Medicine and Lundbeckfonden. H.Y. was funded by the China Scholarship Council. The authors thank the FACS Core Facility at Aarhus University for their help and support.

## Author contributions

H.Y. and D.M.F.v.A conceived the study; H.Y. performed experiments; A.T.F. performed molecular biology;

H.Y. analysed data and H.Y. and D.M.F.v.A. interpreted the data and wrote the manuscript with input from all authors.

## Supplementary figure legends

**Supplementary figure S1: Gating strategy for flow cytometry analysis.** Flow cytometry data were first visualized using forward scatter height (FSC-H) and side scatter height (SSC-H) to gate cells based on size and granularity. Singlets were isolated by comparing FSC-H versus FSC area (FSC-A) and SSC-H versus SSC area (SSC-A). Transfected cells were then identified by gating on mTagBFP2 fluorescence, differentiating mTagBFP2 positive (mTagBFP2+) cells from mTagBFP2 negative (mTagBFP2-) cells. For fixed, permeabilized, and RL2-AF647 stained samples, singlets were visualized on SSC-H versus mTagBFP2 area (mTagBFP2-A) plots to define mTagBFP2+ cells. An untransfected control was used to define the gating threshold. Median fluorescence intensities of OGT-sfGFP, mScarlet3-OGA, and RL2-AF647 were extracted from the mTagBFP2+ population. For samples not requiring fixation and antibody staining, singlets were visualized using RedDot2 area (RedDot2-A) versus mTagBFP2-A to select live, transfected mTagBFP2+ cells. Again, gating was aided by an untransfected control. Median fluorescence intensities of OGT-sfGFP and mScarlet3-OGA were extracted from the mTagBFP2+ population. For samples without plasmid transfection (e.g., OSMI-4 or TMG treatment), singlets were gated based on RedDot2-A to exclude dead cells, and the median fluorescence intensities of OGT-sfGFP and mScarlet3-OGA within the gated singlet population were directly extracted.

For statistical analysis, in transfection experiments, regardless of fixation status, the median fluorescence intensities of OGT-sfGFP, mScarlet3-OGA, and RL2-AF647 (if applicable) from the mTagBFP2+ population were normalized to the corresponding median intensities from the mTagBFP2-population within the same sample. In experiments involving only chemical treatments without transfection, raw median fluorescence intensities were used directly for analysis.

