## Supplementary figures and images for "A triple fluorescence approach to measure O-GlcNAc dyshomeostasis in stem cells"

### Supplementary Figure 1

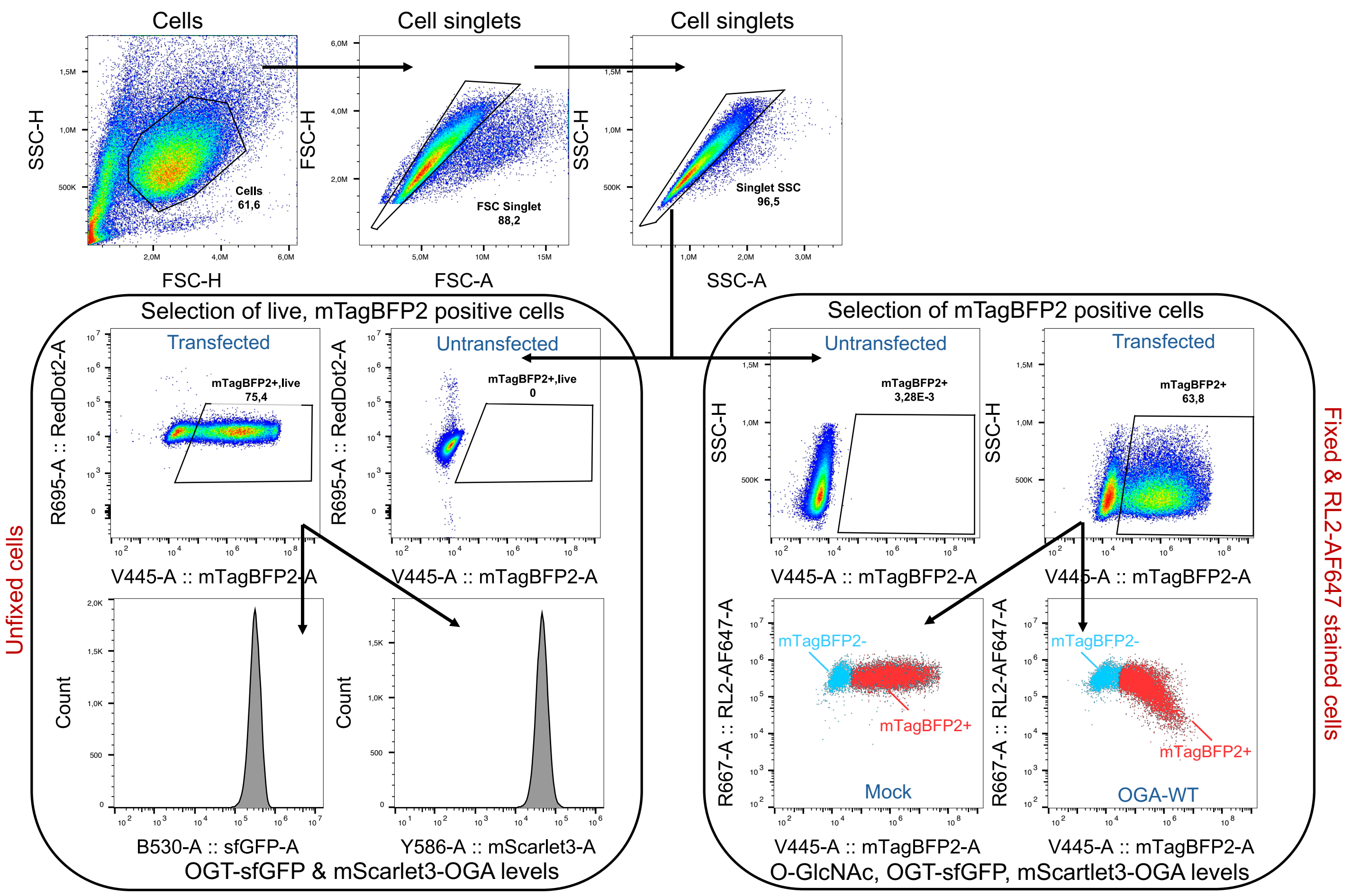
